# Concurrent behavioral modelling and multimodal neuroimaging reveals how feedback affects the performance of decision making in internet gaming disorder

**DOI:** 10.1101/2023.10.22.563399

**Authors:** Xinglin Zeng, Ying Hao Sun, Fei Gao, Lin Hua, Weijie Tan, Zhen Yuan

## Abstract

Internet gaming disorder (IGD) raises questions on how feedback from the previous gaming round affects the risk-taking behavior in the subsequent round. Forty-two participants underwent a sequential risk-taking task, which was measured be behavioral modeling. Concurrent electroencephalogram and functional near-infrared spectroscopy (EEG-fNIRS) recordings were performed to demonstrate when, where and how the previous-round feedback affects the decision making to the next round. We discovered that the IGD illustrated heightened risk-taking propensity as compared to the HCs, indicating by the computational modeling (*p* = 0.028). EEG results also showed significant time window differences in univariate and multivariate pattern analysis between the IGD and HCs after the loss of the game. Further, reduced brain activation in the prefrontal cortex during the task was detected in IGD as compared to that of the control group. Risky decision-making in IGD might be due to the complex interplay between emotional response and other cognitive factors.

## Introduction

Internet Gaming Disorder (IGD) is one kind of representative behavioral addictions, which is characterized as repetitively and compulsively involving internet gaming without considering the adverse consequences (Derevensky, Hayman, & Lynette, 2019). Due to its high prevalence rate (Stevens, Dorstyn, Delfabbro, & King, 2021), IGD is now included in both the 11th revision of the International Classification of Diseases (ICD-11) and the appendix of the 5th edition of the Diagnostic and Statistical Manual of Mental Disorders (DSM-5) (Luo et al., 2021). Similar to the substance use disorder, IGD is due to the dysfunction in dopaminergic neural circuitry engaging in compulsive seeking sensation, the disruption in prefrontal cortex (PFC) involving the executive functions like controlling behaviors, and the deficits in amygdala functions associated with negative emotional reaction (Weinstein & Lejoyeux, 2020). Besides, it is discovered that IGD also exhibits similar impairment in emotion regulation and executive function with other psychiatric disorders, such as major depression disorder, anxiety, attention deficit hyperactivity disorder, and etc. (Ko, Yen, Yen, Chen, & Chen, 2012; Ostinelli et al., 2021). However, in addition to the disrupted executive function and emotion regulation (Argyriou, Davison, & Lee, 2017; Shin, Kim, Kim, & Kim, 2021; Z. Zhou, Zhou, & Zhu, 2016), it is also essential to inspect other complex cognitive process of IGD during task like decision making.

Decision-making is a high-level and dynamic cognitive process in humans, involving weighing alternative outcomes’ desirability and probabilities (Rilling & Sanfey, 2011). According to the criteria for accessing whether the probability of each outcome is predictable, decision making can be classified into two categories: ambiguity and risky decision-making (Y. Li et al., 2019). More importantly, even if the outcome is predictable for risky decision making, the decision-making behaviors are stilled influenced by a bunch of cognitive components such as valence of outcomes (win/lose), amplitude and probability of cost-benefit, characteristics of participants (e.g., risky propensity and inverse temperature), emotional regulation, and etc. (Kusev et al., 2017). Besides, the instantaneous outcome from the previous round of gaming might have a feedback impact on the decision-making patterns of the following round (Figner, Mackinlay, Wilkening, & Weber, 2009; Oskarsson, Van Boven, McClelland, & Hastie, 2009). For example, participants displayed increased risk-seeking behavior following a loss in risk decision-making paradigms (Z. Liu et al., 2020; Xue, Lu, Levin, & Bechara, 2011). Conversely, Pedroni et al. (2017) found that participants exhibited greater risk-taking tendencies after experiencing wins, a phenomenon known as positive recency in sequential decision-making paradigms.

Besides, decision-making behavior is influenced by various factors including loss aversion and risk propensity. As conventional behavioral data analysis may not capture these nuanced characteristics, computational modeling of cognitive tasks has been widely employed in clinical populations to overcome the limitations of traditional behavioral tasks. Previous studies constructed computational models to reflect individual learning ability, sensitivity to wins and losses, inverse temperature, and risk-taking propensity in sequential risky decision making such as the Balloon Analogue Risk Task (BART) (H. Park, Yang, Vassileva, & Ahn, 2021; van Ravenzwaaij, Dutilh, & Wagenmakers, 2011; R. Zhou, Myung, & Pitt, 2021).

Existing findings on the behaviors of IGD in risky decision-making were rather mixed (Y.-W. Yao, Zhang, Fang, Liu, & Potenza, 2022). Some studies reported no group difference of risky propensity between individuals with IGD and health controls (HCs) (Deleuze et al., 2017; Y. W. Yao, Chen, et al., 2015). By contrast, individuals with IGD were also found to exhibit reduced loss aversion and make riskier choices in the loss domain (Y. W. Yao, Chen, et al., 2015), while participants with IGD made riskier choices in the gain domain (Ko et al., 2017). These findings suggested that the risky propensity of individuals with IGD may be modulated by some confounding factors. Specifically, no studies have explored whether IGD manifests different behavioral patterns influenced by the valence of the previous trial (win or loss outcome). Thus, to better evaluate the affecting factors in risky propensity, this study aims to examine the effects of outcome valence on subsequent decision-making performance in individuals with IGD using a sequential risk-taking paradigm (Brassen, Gamer, Peters, Gluth, & Büchel, 2012).

Furthermore, to investigate the potential neural mechanisms of IGD in sequential risky decision-making, we will employ functional near-infrared spectroscopy (fNIRS) neuroimaging and electrophysiology (EEG) simultaneously. The combination of EEG and fNIRS would provide valuable insights into the neural dynamics underlying decision-making, as they capture event-related potentials (ERPs), brain oscillation characteristics, and oxygenated hemoglobin (HbO) concentration changes of perceiving outcomes, during decision making process (Fei Gao et al., 2023; F. Gao et al., 2022; R. Li et al., 2022). Previous studies have reported abnormal activation in the prefrontal cortex (PFC) of individuals with IGD using functional Magnetic Resonance Imaging and the BART (L. Liu et al., 2017; Qi et al., 2015), Thus, fNIRS will be used to measure changes in HbO concentration in the PFC during sequential risky decision-making. In addition, studies utilizing brain electrophysiological activities have found higher brain oscillations in slow-wave activity (theta, beta) among individuals with IGD (Y. J. Kim et al., 2017) and observed differences in feedback-related negativity (FRN) and P300 amplitudes in risk decision-making tasks (Q. Li et al., 2020; Yau, Potenza, Mayes, & Crowley, 2015; Zhao, Li, Hu, Wu, & Liu, 2017). Especially, compared to univariate pattern analysis (ERPs), the multi-variates pattern analysis (MVPA) can decode neural activities across multiple EEG channels, with enhanced sensitivity, integrated information, reduced the noise influence, etc (Grootswagers, Wardle, & Carlson, 2017). In light of the MVPA strength and no previous studies have adopted this technique to decode the neural activity of perceiving feedbacks, we wish to explore the different time course of potential neural activity perceiving the valence of feedbacks among IGD and HCs.

To date, it is yet unclear whether the win/loss outcome in feedback from the previous round of gaming will promote or inhibit the risk-taking behaviors during the next round. In particular, the neural mechanism on how the win or loss outcomes affect further decision-making performance during prolonged continuous gaming is also unclear. Therefore, this aim of this study is to inspect the underlying cognitive neural mechanism of risky decision-making in IGD, demonstrating the impact of previous gaming outcomes (feedback) on further decision-making performance. It is hypothesized that IGD patients who lose the previous round of gaming tend to take riskier behaviors in performing the following decision making task as compared to HCs. To test the hypothesis, concurrent behavioral modeling and electroencephalogram-functional near-infrared spectroscopy (EEG-fNIRS) multimodal neuroimaging were carried out to systematically characterize the specific impairments in IGD with a sequential risk-taking task. By using the computational model, nuanced cognitive processes during the task were able to be detected, revealing the aberrant risky decision-making performance. In particular, the developed model was able to describe to what extent the adjustment of behaviors in further decision making is affected by the outcomes of previous trial. Meanwhile, EEG-fNIRS multimodal recordings aid to reveal when and where decision was made in the cortex under the influence of previous gaming results. It is expected that the present study might provide new insights for understanding the cognitive neural mechanism associated with risky decision-making in IGD.

## 2 Method

### 2.1 Participants

Forty-two right-handed participants including 15 individuals with IGD (mean age: mean age: 21.46, 6 females) and 27 health controls (HC, mean age: 21.67, 17 females) were recruited from the University of Macau. The inclusion criteria for IGD were as follows: 1) aged between 20 and 30 years with more than 12 years of education; 2) played internet games for at least 4 hours per day on weekdays and 8 hours per day on weekends, or a total of 40 hours per week, for a minimum duration of 2 years, as confirmed by total game time records. Participants in the control group were gender-, age-, and education-level matched with those in the IGD group and had a nonessential internet use of less than 4 hours per day in their daily lives. A semi-structured interview was conducted to assess the DSM-5 criteria for IGD, and participants needed to fulfill at least five out of nine items as defined by the DSM-5 definition (Luo et al., 2021). All individuals with IGD completed the Chen Internet Addiction Scale (CIAS) (Chen, Weng, Su, Wu, & Yang, 2003). Together, CIAS and DSM-5 were used to indicate the severity of internet addiction. The Barratt Impulsiveness Scale (BIS-11), a widely used assessment of impulsiveness, was administered to all participants (X.-y. Li et al., 2011). All participants signed informed consent forms prior to the experiment. The protocol and all procedures of this study were approved by the Institutional Review Board with the University of Macau.

### 2.2 Procedure

Participants were seated in a quiet room, in which both behavioral and EEG-fNIRS data were collected concurrently during the task. Before the formal experiment, participants were informed to try to receive the rewards as more as possible during the task because their payments were decided by their rewards. Initially participants were required to take the practice test with 16 trials to ensure they were familiar with the experimental procedure. And then participants performed the formal test with 200 trials, which took about 30-40 minutes.

Each trial started with a fixation in the center of the monitor ranged from 1 to 3 seconds, followed by a decision-making task with 1.5-5.5 seconds, a feedback task of 3 seconds, and a rating period of 3 seconds (Fig. 1). At the beginning of decision-making task, eight closed boxes were presented in the screen of the PC, in which 7 boxes contained gains (gold) or 1 containing a loss (devil) was randomly assigned to one of the eight boxes. And then the boxes were opened from the left to the right sequentially. When one box was opened, participants needed decide either opening the next box or stopping to collect all the gains acquired so far in that trial by pressing the keyboard in 2000ms, or the next boxes would be opened automatically. The decision-making period task was over when the devil box was opened (defined as lose condition) or when participants decided stopping to collect all the gains (defined as gain condition) before the next box was opened. The feedback task lasted 3000 ms, demonstrating that participants either won or lost the game with different colors of box. The screen also displayed the actual position of the devil, thus informing participants how many points they’d collected or how many chances they’d missed at the same time. Finally, participants were required to rate the feelings about their choice of this trial within 3000 ms by using a 9-point scale from extreme regret (defined as −4) to extreme relief (defined as 4).

**Fig. 1.**
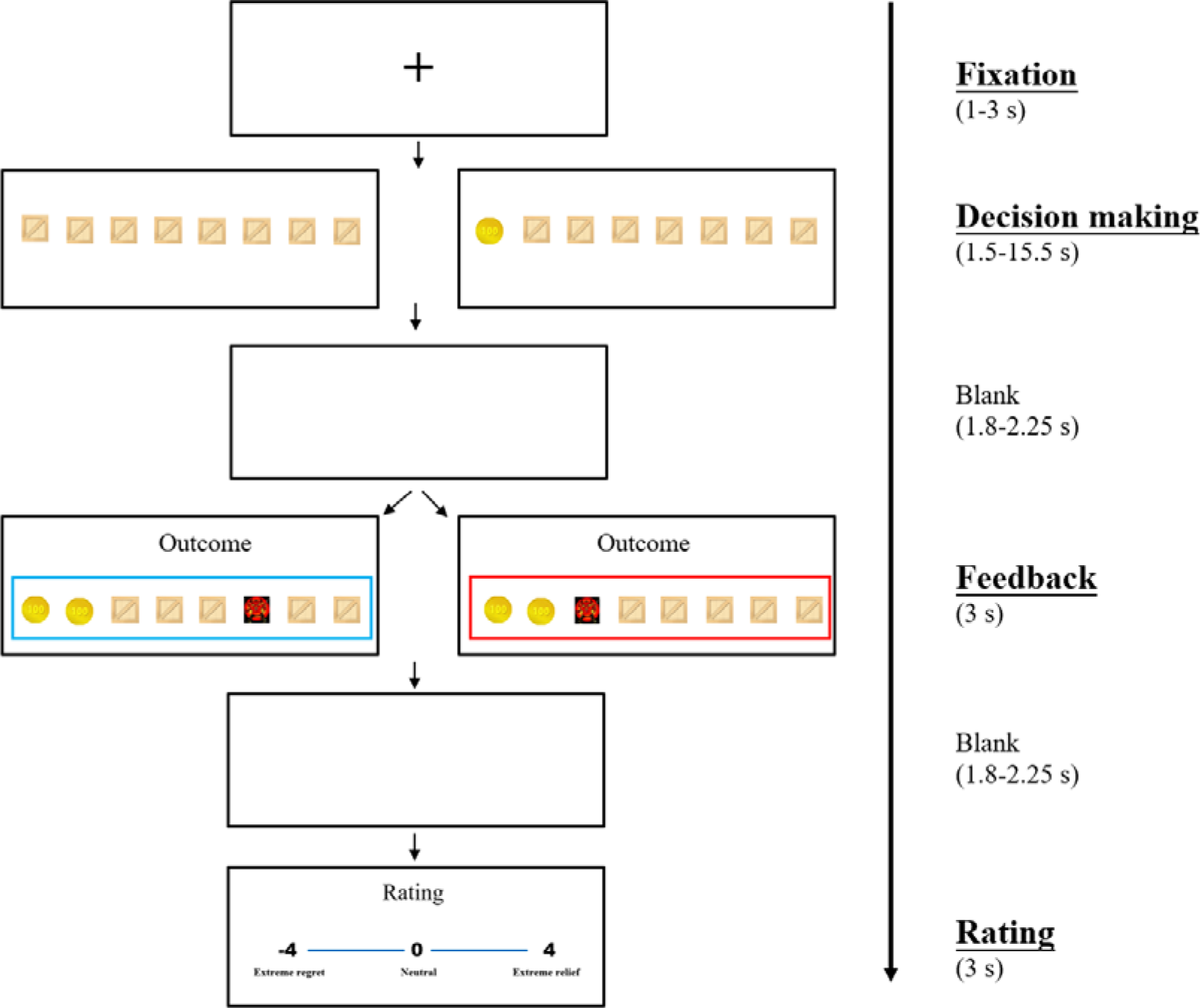
Schematic of the sequential risk-taking task. The trial of sequential risk-taking task was composed of fixation period, decision making period, feedback presentation period and rating period.

### 2.3 Behavioral modelling

#### 2.3.1 Model details

As the procedure mentioned earlier, the probability of meeting devils (losing the coins of that trial) 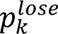 is 1/8. Thus, according to van Ravenzwaaij et al. (2011), the expected utility after l boxes on trial k, *U_kl_* can be presented as:

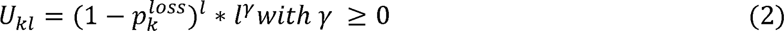

the optimal boxes opened (*v_k_*) can be calculated by setting the first derivative of Eq. (4) for *l* equals zero. Here we can get the optimal boxes opened:

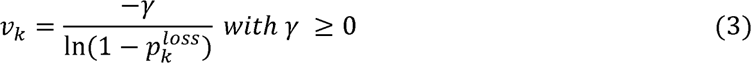

Considering the randomness trait of decision making, here we use the parameter τ, to present inverse temperature of participants, in which higher τ value indicates more deterministic behavior of the decision making. So, we could calculate the probability 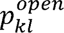 if the participants opened the boxes *l* on trial k.

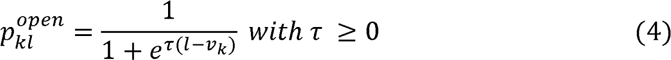

Considering that the behaviors in the next trial would be influenced by the emotional response of previous trial, here we proposed a revised model (parameter-3 model). In this model, we used a parameter *a* to present the degree of the participants being influenced by emotional response of last trial. Thus, the probability 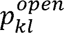, can be rewritten as:

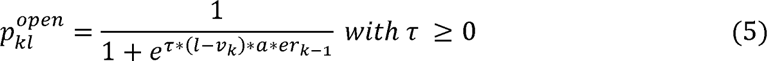

The *er_k_*_-1_is the emotional rating scores in last trial.

#### 2.3.2 Parameter Estimation

The parameters of the two computational models were estimated through a hierarchical Bayesian analysis framework, as outlined by Lee (2011). This approach enables the simultaneous estimation of both individual and group-level data in a mutually constraining manner. The implementation of hierarchical Bayesian analysis utilized the Stan software package (https://mc-stan.org/) and the hBayesDM package (Ahn, Haines, & Zhang, 2017) within the R programming environment (http://www.r-project.org/). We employed the Hamiltonian Monte Carlo method to facilitate the estimation process. To ensure convergence to the desired distributions, a substantial sample size of 4000 was employed. Additionally, we ran four independent chains to assess whether the posterior distributions remained independent of the initial starting points. This rigorous approach was designed to enhance the robustness and reliability of our parameter estimates.

#### 2.3.3 Model Comparison

The performance of the two models was compared using the leave-one-out information criterion (LOOIC) (Vehtari, Gelman, & Gabry, 2017). LOOIC was calculated based on leave-one-out cross-validation, estimating out-of-sample prediction accuracy from the log-likelihood derived from the posterior distributions. A lower LOOIC score indicates a better model performance. LOOIC weights, defined by Akaike weights (Wagenmakers, Farrell, & review, 2004) based on LOOIC values, were used to compare the models.

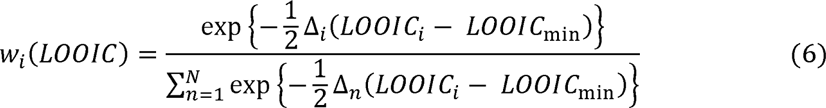

### 2.4 Concurrent fNIRS and EEG recordings

The fNIRS-EEG data acquisition cap (EasyCap, Herrsching, Germany) with reference to the international 10-20 system was used for concurrent fNIRS and EEG recordings. The fNIRS data were collected by NIRScout system (NIRx Medizintechinik GmbH, Berlin, Germany) with 8 LED sources and 8 optical detectors, yielding 20 measurement channels with an inter-optode distance of 30 mm placed along the frontal regions. The light attenuation was measured at the wavelength of 760 nm and 850 nm with the sampling rate of 7.81 Hz. In addition, the Montreal Neurological Institute (MNI) coordinates of each fNIRS channel were quantified using ICBM-152 head model (Singh, Okamoto, Dan, Jurcak, & Dan, 2005) and the NIRSite 2.0 toolbox (NIRx Medizintechnik GmbH, Berlin, Germany). The coordinates were then imported into the NIRS-SPM software toolbox for spatial registration (Ye, Tak, Jang, Jung, & Jang, 2009). The spatial configurations (MNI coordinates, Brodmann area, anatomical label, and percentage of overlap) of the 20 channels were provided in Fig. 2 and Table 1.

**Fig. 2.**
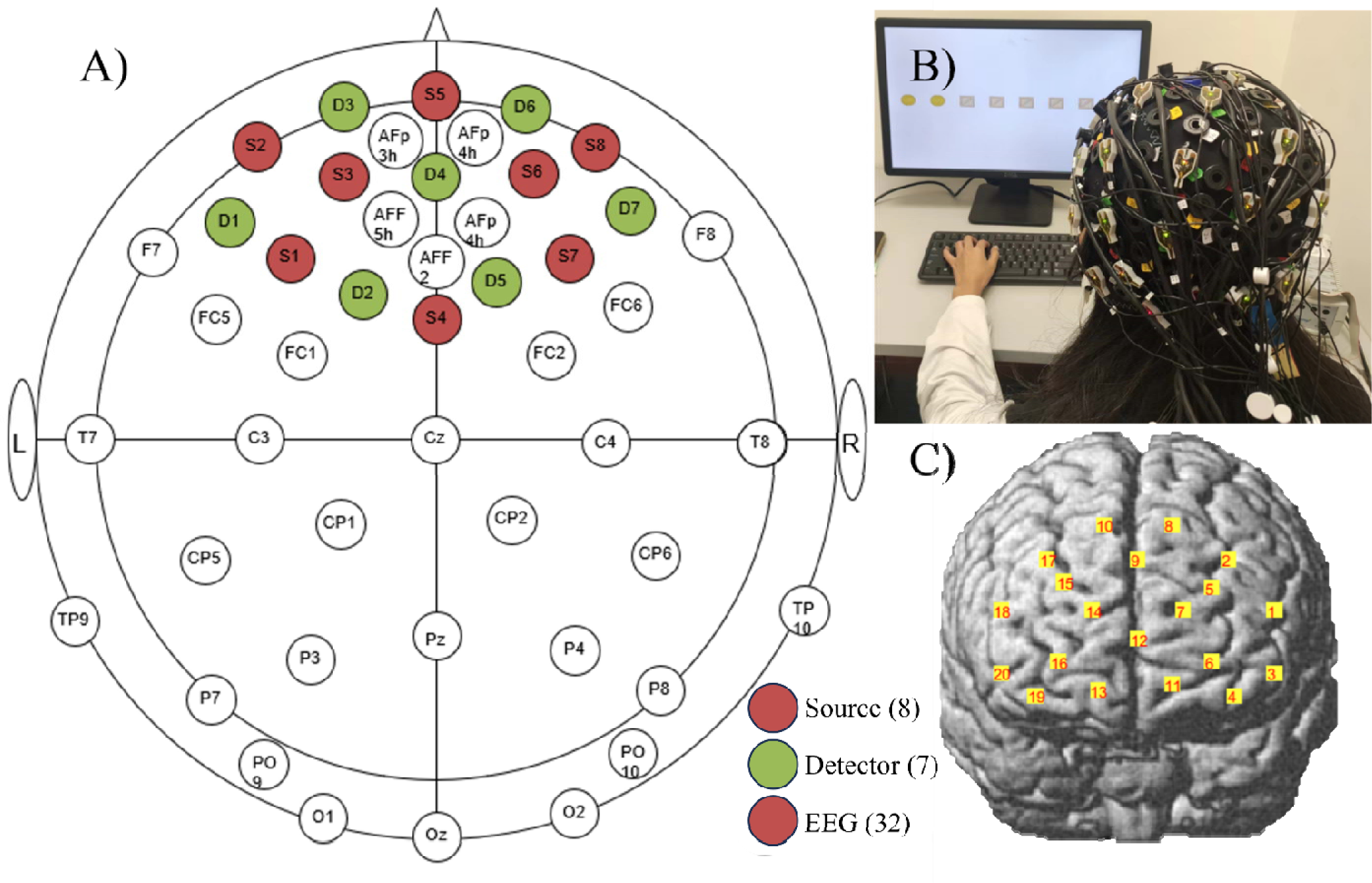
The fNIRS-EEG device and location. A) The fNIRS channel and EEG location; B) The fNIRS-EEG device setup; C) fNIRS channels reconstructed by NIRS-SPM.

**Table 1.**
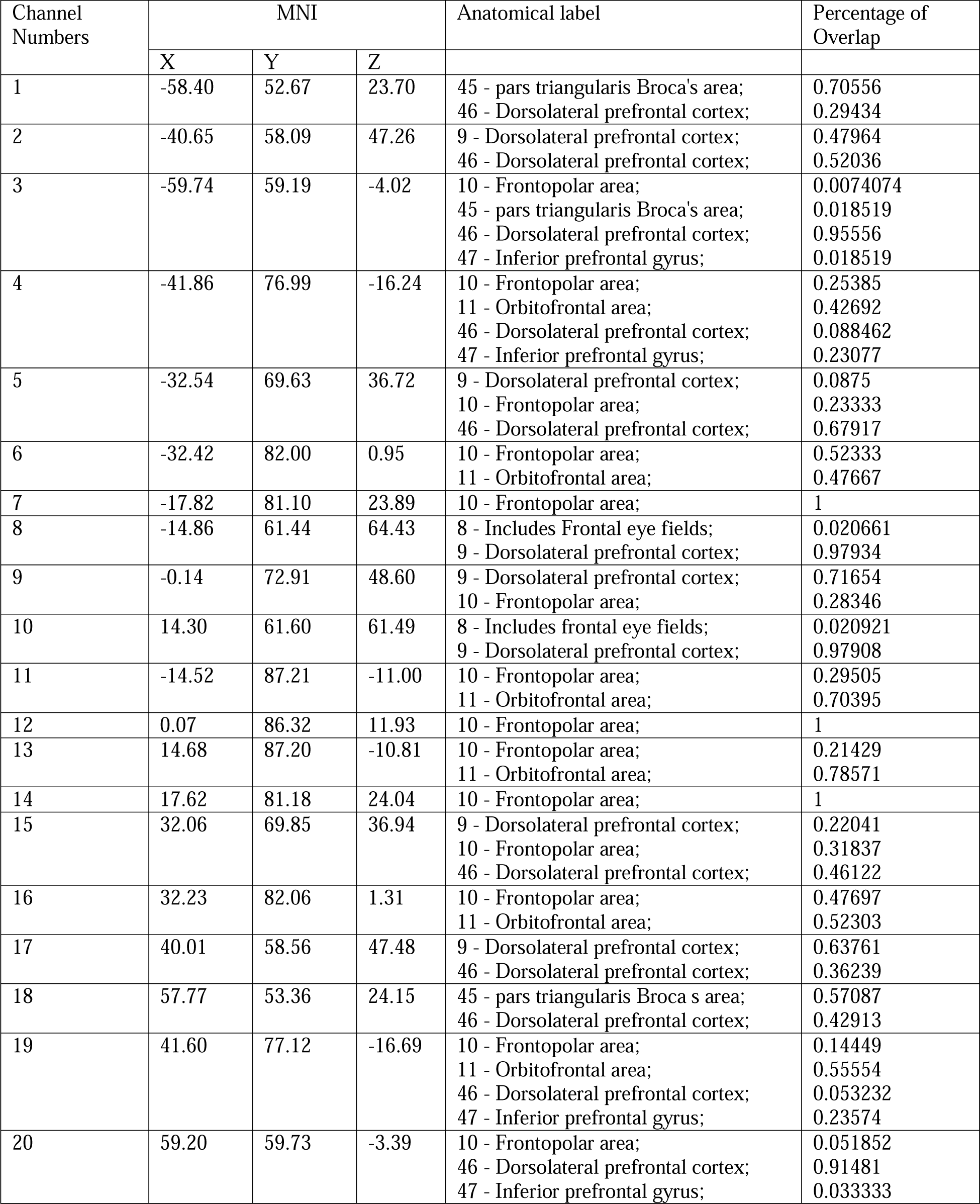
3D MNI coordinates, anatomical labels, and coverage percentage of fNIRS channels.

| Channel Numbers | MNI |  |  | Anatomical label | Percentage of Overlap |
| --- | --- | --- | --- | --- | --- |
|  | X | Y | Z |  |  |
| 1 | -58.40 | 52.67 | 23.70 | 45 - pars triangularis Broca's area;<br>46 - Dorsolateral prefrontal cortex; | 0.70556<br>0.29434 |
| 2 | -40.65 | 58.09 | 47.26 | 9 - Dorsolateral prefrontal cortex;<br>46 - Dorsolateral prefrontal cortex; | 0.47964<br>0.52036 |
| 3 | -59.74 | 59.19 | -4.02 | 10 - Frontopolar area;<br>45 - pars triangularis Broca's area;<br>46 - Dorsolateral prefrontal cortex;<br>47 - Inferior prefrontal gyrus; | 0.0074074<br>0.018519<br>0.95556<br>0.018519 |
| 4 | -41.86 | 76.99 | -16.24 | 10 - Frontopolar area;<br>11 - Orbitofrontal area;<br>46 - Dorsolateral prefrontal cortex;<br>47 - Inferior prefrontal gyrus; | 0.25385<br>0.42692<br>0.088462<br>0.23077 |
| 5 | -32.54 | 69.63 | 36.72 | 9 - Dorsolateral prefrontal cortex;<br>10 - Frontopolar area;<br>46 - Dorsolateral prefrontal cortex; | 0.0875<br>0.23333<br>0.67917 |
| 6 | -32.42 | 82.00 | 0.95 | 10 - Frontopolar area;<br>11 - Orbitofrontal area; | 0.52333<br>0.47667 |
| 7 | -17.82 | 81.10 | 23.89 | 10 - Frontopolar area; | 1 |
| 8 | -14.86 | 61.44 | 64.43 | 8 - Includes Frontal eye fields;<br>9 - Dorsolateral prefrontal cortex; | 0.020661<br>0.97934 |
| 9 | -0.14 | 72.91 | 48.60 | 9 - Dorsolateral prefrontal cortex;<br>10 - Frontopolar area; | 0.71654<br>0.28346 |
| 10 | 14.30 | 61.60 | 61.49 | 8 - Includes frontal eye fields;<br>9 - Dorsolateral prefrontal cortex; | 0.020921<br>0.97908 |
| 11 | -14.52 | 87.21 | -11.00 | 10 - Frontopolar area;<br>11 - Orbitofrontal area; | 0.29505<br>0.70395 |
| 12 | 0.07 | 86.32 | 11.93 | 10 - Frontopolar area; | 1 |
| 13 | 14.68 | 87.20 | -10.81 | 10 - Frontopolar area;<br>11 - Orbitofrontal area; | 0.21429<br>0.78571 |
| 14 | 17.62 | 81.18 | 24.04 | 10 - Frontopolar area; | 1 |
| 15 | 32.06 | 69.85 | 36.94 | 9 - Dorsolateral prefrontal cortex;<br>10 - Frontopolar area;<br>46 - Dorsolateral prefrontal cortex; | 0.22041<br>0.31837<br>0.46122 |
| 16 | 32.23 | 82.06 | 1.31 | 10 - Frontopolar area;<br>11 - Orbitofrontal area; | 0.47697<br>0.52303 |
| 17 | 40.01 | 58.56 | 47.48 | 9 - Dorsolateral prefrontal cortex;<br>46 - Dorsolateral prefrontal cortex; | 0.63761<br>0.36239 |
| 18 | 57.77 | 53.36 | 24.15 | 45 - pars triangularis Broca's area;<br>46 - Dorsolateral prefrontal cortex; | 0.57087<br>0.42913 |
| 19 | 41.60 | 77.12 | -16.69 | 10 - Frontopolar area;<br>11 - Orbitofrontal area;<br>46 - Dorsolateral prefrontal cortex;<br>47 - Inferior prefrontal gyrus; | 0.14449<br>0.55554<br>0.053232<br>0.23574 |
| 20 | 59.20 | 59.73 | -3.39 | 10 - Frontopolar area;<br>46 - Dorsolateral prefrontal cortex;<br>47 - Inferior prefrontal gyrus; | 0.051852<br>0.91481<br>0.033333 |

EEG data were recorded with 32 Ag/AgC1 active electrodes (Brain Products, Munich, Germany) attached to the EEG cap (sampling rate: 500 Hz, bandpass: 0.03-70 Hz; reference: Cz). Five EEG electrodes were re-located to their neighboring channels due to the pre-emption of fNIRS probes (Fig. 2A). The electrical impedance of each channel was maintained below 25 kΩ before concurrent data acquisition.

### 2.5 EEG data processing

#### 2.5.1 Data Pre-processing

The EEG data were preprocessed using Matlab R2016a (MathWorks, Natick, USA) and EEGLAB, with a re-reference to grand average and a bandpass filter of 1-30 Hz. Independent Component Analysis (ICA) was applied to correct artifacts due to eye movements (blinks and shifts), muscle activity, and cardiac interference. Channels with high impedance were interpolated using the spherical interpolation method. Subsequently, the clean EEG data were segmented from 100 ms before the onset of the outcome to 1000 ms after the onset. Baseline correction was applied to all epochs, using the mean voltage over the 100 ms preceding the outcome onset as the reference. Epochs with EEG voltages exceeding a threshold of ±75 μV were excluded from further analysis. Two participants from the HC group were excluded due to the large noise of EEG signals.

#### 2.5.2 Global field power (GFP) and event-related potential (ERP) analysis

GFP was used to assess the strength of the electric field at the scalp. GFP is computed as the square root of the average of the squared voltage values recorded at each electrode, serving as an index for the spatial standard deviation of the electric field at the scalp (Murray, Brunet, & Michel, 2008). A larger GFP value signifies a stronger electric field. Differences in GFP waveform data were examined in relation to time, particularlly when the post-stimulus data were significantly deviated from the baseline across various conditions. The significance of GFP amplitude was determined by its consecutive exceedance of a 95% confidence interval for at least 20 ms relative to the 100-ms pre-stimulus baseline (Hua, Gao, Leong, & Yuan, 2023). Subsequently, GFP peaks were identified based on the averaged GFP waveform across participants.

To identify the topographic modulations, global map dissimilarity measures (GMD) were carried out by using randomization statistics. GMD is calculated as the root mean square of the difference between strength-normalized vectors (Murray et al., 2008). Differences in GMD values were also analyzed with respect to time. The statistical significance of GMD was demonstrated by using a topographic analysis of variance (ANOVA) with 10,000 permutations (p < 0.05). Importantly, GMD is independent of field strength, whereas the significant GMD results indicate the differences in neural generators across various conditions. Significance levels of GMD results were adjusted by using a duration threshold of 20 ms.

For the present study, electrodes located in the central regions were selected for analysis: FC1, FC2, C3, C4, CP1, and CP2. All the trials for each participant were averaged and then paired two-sided t-tests and independent two-sided t-tests were performed to detect the significant differences in relation to feedback outcomes and group. The *p* values were corrected using the false discovery rate (FDR) with a significance level of *p* = 0.05.

#### 2.5.3 Time-frequency analysis

To inspect the potential brain oscillations of ERPs, MNE Python software 1.2 was utilized to compute event-related oscillations (EROs). Morlet wavelets were applied to extract the time–frequency representation of induced power for a given time point (550 data points, ranged from −100 ms to 1000 ms, relative to feedback onset) and frequency (4-30 Hz with a resolution of 2 Hz) (Morales & Bowers, 2022). The Morlet wavelet transform is generally used to generate EEG spectral components as it strikes a favorable balance between temporal and frequency resolution. More specifically, Morlet wavelet transform demonstrates its exceptional efficiency when applied to non-stationary signals, making it an excellent candidate to analyze the present data (Vecchio et al., 2022). Statistical analysis of EROs was performed with a non-parametric procedure based on permutations (1000 times) and cluster-level statistics (*p* = 0.001, cluster-level *p* = 0.01).

#### 2.5.4 Multivariate pattern analysis

MVPA (MNE Python software version 1.2) was used to examine the topographic weighting of EEG signals that was able to effectively differentiate the win and loss feedback within specific time intervals. A linear classifier based on L2-regularized logistic regression was used to identify optimal projections of the sensor space, enabling the discrimination between win and loss feedback in individuals at specific time points. This approach allowed for the investigation of feedback perception availability in IGDs and HC groups based on stimulus-locked EEG data, and the temporal dynamics of this availability were accessed by using time-resolved decoding.

The accuracy of the linear L2-regularized logistic regression classifier, trained on multi-electrode single-trial EEG signals, was independently assessed for each time point. A Monte Carlo cross-validation procedure (n = 100) was repeated for 10 times, with the entire dataset randomly partitioned into 10 subsets consisting of a training set (90% of the trials) and a test set (the remaining 10%). These time windows corresponded to the ERP and ERO analyses, spanning from 100 ms before the stimulus onset to 1000 ms after the stimulus onset. Individual classification accuracy was considered statistically above chance when it exceeded the classification accuracy obtained from permuted labels (paired t-test, α = 0.05). Group-level analysis followed the same procedure, with group averages computing the across individual averages. Correction for multiple comparisons was achieved by using a time-cluster-based approach, in which a time point was considered significant only if it belonged to a cluster comprising at least twenty consecutive significant time points.

### 2.6 fNIRS data analysis

fNIRS signals from five participants (three in IGD and two in HC groups) were not correctly recorded due to machine malfunction, and data from one HC was excluded for further analysis due to the large noise. fNIRS data were preprocessed by using the nirsLAB toolbox (NIRx Medizintechnik GmbH, Berlin, Germany). Motion artifacts were removed and detrending was applied to the raw data by using the built-in algorithm. To achieve the best signal-to-noise ratio, the data were further band-pass filtered with a low cutoff frequency of 0.1 Hz and a high cutoff frequency of 0.01 Hz to minimize physiological noise associated with heart pulsation (1-1.5 Hz), respiration (0.2-0.5 Hz), and blood pressure waves (∼0.1 Hz) (Lu et al., 2021). The HbO changes were modeled by using statistical parametric mapping with the canonical hemodynamic response function, resulting in general linear model coefficients (beta values) across all conditions for each participant. In this study, only HbO signals were analyzed as they provide a more sensitive indicator of changes associated with regional cerebral blood flow (Xu et al., 2020). Subsequently, the beta values of feedback perception in each channel were compared using a two-way ANOVA and corrected using FDR.

### 2.7 Statistics analysis

Following a case-control study protocol, the statistical analyses were conducted using R software 3.6 and MATLAB 2016. Risk-taking behavior was defined as the number of opened win boxes in the gain condition, which reflects a standard measure in sequential risk-taking tasks (Z. Liu et al., 2020). Two-way ANOVA analysis was used to explore the factors (participants and outcome of the last trial) that affected risk behaviors in the subsequent trial. Continuous demographic data (age, DSM-V, CIAS, BIS-11) were compared using independent t-tests, and gender distribution between the two groups was compared using a chi-squared test. The statistics for EEG and fNIRS analyses are described in each respective section. The parameters of computational models were compared using independent t-tests. Potential correlations between demographic data (CIAS, BIS-11) and modeling parameters, as well as the number of boxes opened in the gain condition, were explored using Pearson correlation.

## 3 Results

### 3.1 Demographic data

There were no significant differences in age and gender between the IGD and HC groups (Table 2). However, significant differences were detected in education levels, DSM-V scores, CIAS score, and BIS-11 scores between the IGD and HC groups.

**Table 2.** Demographic and personality characteristics of the included participants.

|  | **IGD (15)**<br>**Mean (std)** | **HC (27)**<br>**Mean (std)** | ***t* value** | ***p* value** |
| --- | --- | --- | --- | --- |
| **Age** | 21.4667 (1.993) | 21.667 (2.019) | -0.3252 | 0.7472 |
| **Gender** | 6 female / 9 male | 17 female /10 male | $X^2=1.23$ | 0.2674 |
| **Education** | 14.4667 (1.1254) | 15.481 (1.889) | -2.181 | 0.0352 |
| **DSM-V** | 6.6 (0.8281) | 1.926 (0.8738) | 17.183 | <0.001 |
| **BIS-11** | 61.533 (5.0124) | 47.481 (5.366) | 8.4867 | <0.001 |
| **CIAS** | 77.20 (7.013) | 40.037 (6.531) | 16.861 | <0.001 |

### 3.2 Behavioral results

Behavioral results demonstrated that individuals with IGD tended to open more boxes as compared to the HC group (*F* (2, 40) = 12.397, *p* = 0.0007, Fig 3 Panel A). Interestingly, both the IGD and HC groups showed a greater likelihood of opening more boxes after loss outcomes as compared to that of win outcomes, as indicated by paired t-tests (IGD: boxes opened _Gain_ = 3.914 ± 0.763, boxes opened Lose = 4.111 ± 0.737, *t* = −2.550, *p* = 0.03; HC: boxes opened Gain = 3.402 ± 0.574, boxes opened Lose = 3.620 ± 0.519, *t* = −3.135, *p* = 0.004).

**Fig. 3.**
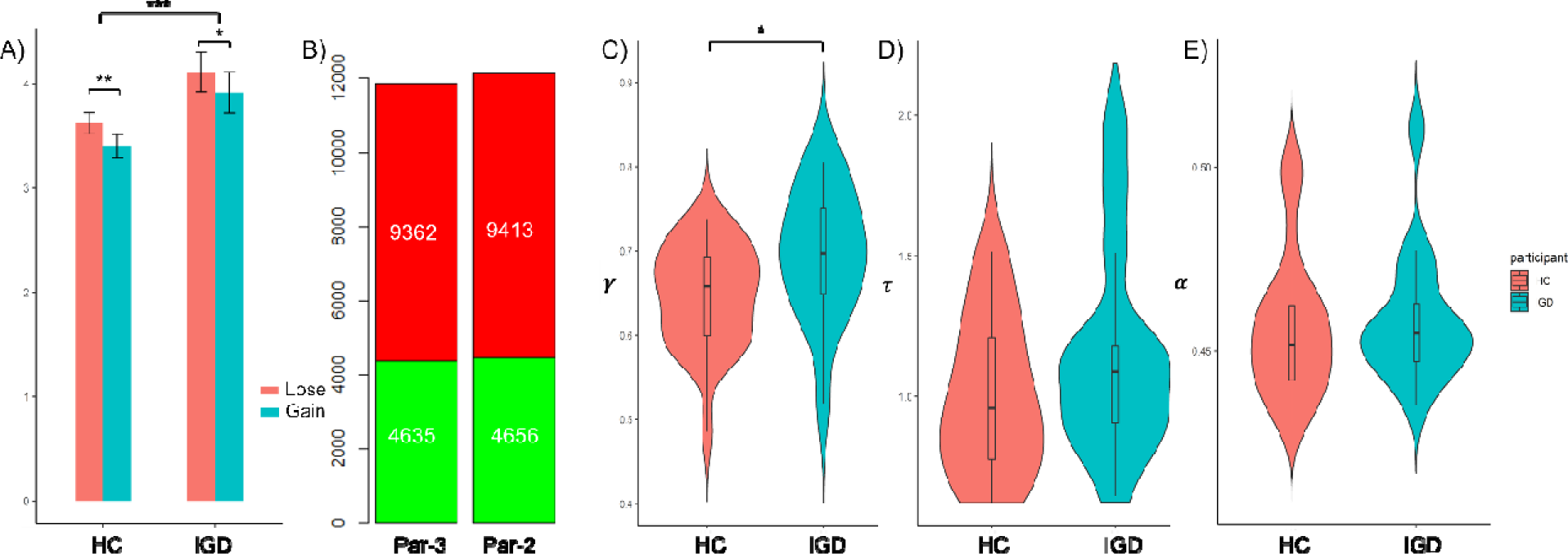
The behavioral and modeling results. **A).** The significant differences in the opened box number between the gain and lose conditions and between the HC and IGD groups; B). The LOOIC values calculated from the tw models; C). The computational modeling parameter rho, indicating risk taking propensity; D). The computational modeling parameter tau, indicating inverse temperature; and E). The computational modeling parameter alpha, indicating emotional affection. *** indicates p<0.001; ** indicates p<0.01; while * indicates *p*<0.05.

**Fig. 4.**
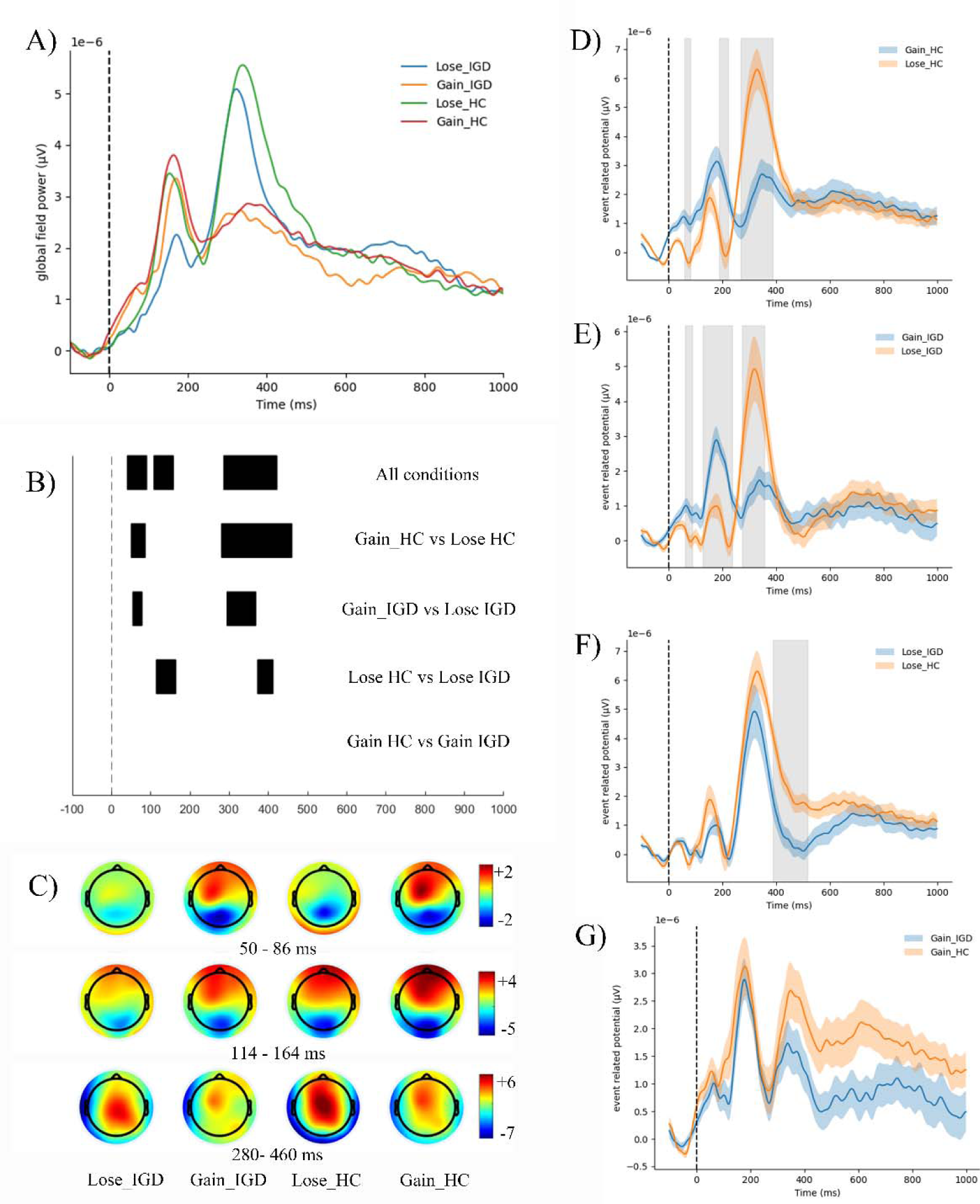
Neural responses to loss/win feedbacks exhibited significant difference in GFP and ERP between the IGD and HC groups. **A).** GFP to loss/win feedbacks for both the IGD and HC groups; **B).** The time segments of significant GMD were indicated in black bars. The top row depicts the significant main difference of the four conditions, whereas the other rows depict the pairwise comparison (HC_Gain_ vs HC_Lose_, IGD_Gain_ vs IGD_Lose_, HC_Gain_ vs IGD_Gain_, HC_Lose_ vs IGD_Lose_); **C).** Topography of the four conditions regarding the three time windows (50-86 ms, 114-164 ms, and 280-460 ms). For each topography, EEG signals were averaged across all time points within specific time window. Color bar denotes the voltage value (μV).; **D-F).** The ERP pair wise comparison to loss/win feedbacks for both the IGD and HC groups. Colored shaded areas: mean ± error bar; light gray shaded areas: the period of significant decoding at the group level (p_FDR_<0.05).

**Fig. 5.**
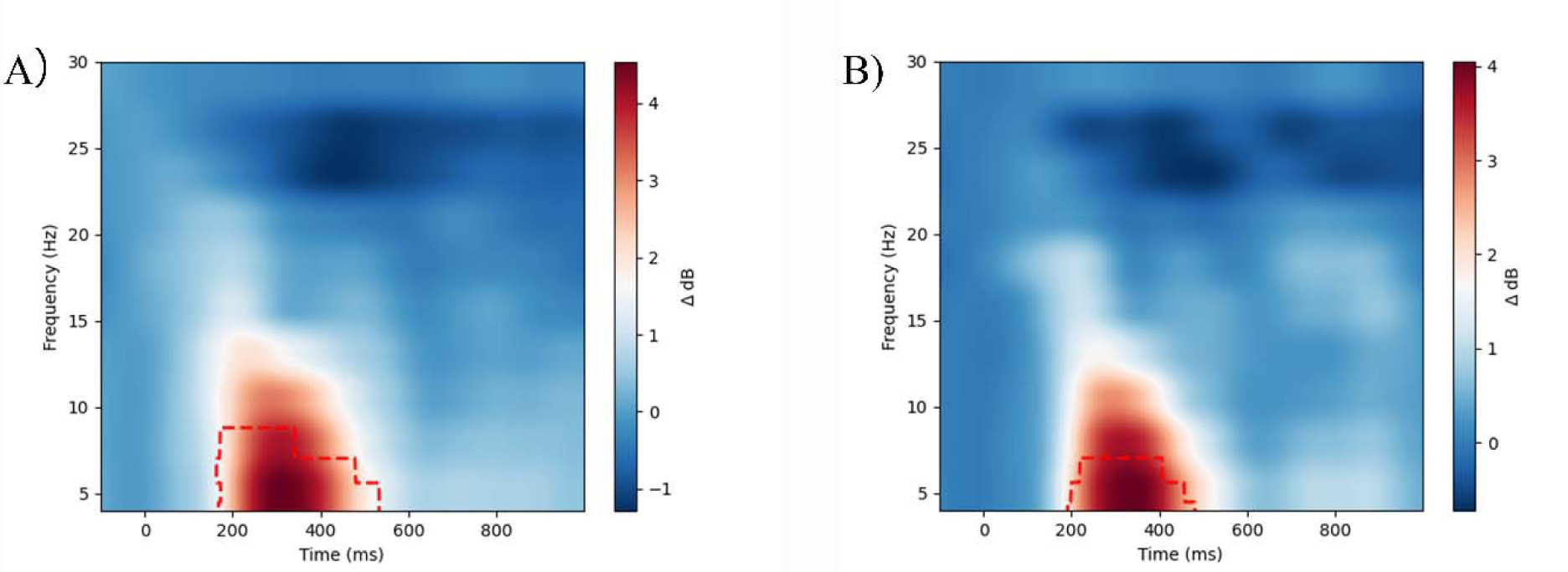
EROs to feedbacks exhibited significant difference. A) EROs to lose feedback, contrasting to gain feedbac of EROs in the HC groups; B) EROs to lose feedback, contrasting to gain feedback of EROs in the IGD group.

### 3.3 Modeling results

The results comparison using the leave-one-out information criterion (LOOIC) demonstrated that the Par-3 model involving the impact of emotional influence, provided a better fit to the data as compared to the Par-2 model (Fig 3 Panel B).

The posterior distributions of fitted Par-3 model parameters were calculated to quantify the underlying cognitive processes during the sequential risk-taking task. The IGD group exhibited a significantly higher risk-taking propensity (IGD = 0.697 ± 0.058) as compared to the HC group (HC = 0.644 ± 0.077, *t* = 2.339, *p* = 0.0284, Fig 3 Panel C). However, no significant differences were detected between the two groups in terms of behavioral consistency (τ_IGD_ = 1.295, τ_HC_ = 1.165, *t* = 0.374, *p* = 0.7117, Fig 3 Panel D) or emotional affection (a_IGD_ = 0.484, a_HC_ = 0.481, *t* = 0.631, *p* = 0.533, Fig 3 Panel E).

### 3.4 GFP and ERP results

Both the IGD and HC groups exhibited significantly different neural responses to loss and win feedbacks from 50 ms to 460 ms. Interestingly, three GFP peaks with time periods between 50-86 ms, 114-164 ms and 280-460 ms showed significant difference between the four conditions, corresponding to ERP component P1, feedback-related negativity (FRN), and P3, respectively. In particular, for both the HC and IGD groups, the win feedbacks induced higher electric field strength in the 50-86 ms time window (higher positive potential in the central region and higher negative potential in the occipital region), and lower electric field strength in the 280-460 ms time window (lower positive potential in the central region), as compared to loss feedbacks. In addition, the loss feedback in IGD group induced significant differences in electric field strength (lower positive potential in the prefrontal region and lower negative potential in the occipital region) during the time period 114-164 ms time as compared to that in HC group. Further, no significant differences in the topographical distribution of electric field, independent of the electric field strength, were detected after win feedbacks between the IGD and HC groups.

Interestingly, for both IGD and HC groups, the loss feedbacks elicited a higher P1 amplitude, lower FRN amplitude, and higher P300 amplitude as compared to win ones. Besides, the IGD group demonstrated a shorter P300 latency compared to the HC group.

### 3.5 ERO results

Significant differences in frequency spectra between the win and loss feedbacks were detected for both the IGD and HC groups. In HC group, delta power of EROs after loss feedbacks was significantly higher in the 174-532 ms time window compared to those of win feedbacks. Likewise, for the IGD group, higher delta power of EROs was identified in the 192-456 ms time window. More importantly, no significant differences in frequency spectra after win or loss feedbacks were detected in either the IGD or HC group.

### 3.6 One-vs-one decoding results

To further inspect how the global perceived orientation varied across the four conditions, one-vs-one classification was performed for all paired comparisons. Pairwise classifications yielded significant accuracies above chance level at specific time points across the trials (*p* < 0.05, lasting for > 20 ms). In the HC group, pronounced decoding performance was detected between win and loss feedbacks, roughly from 140 to 570 ms, indicating a long-lasting neural response difference. In the IGD group, the decoding performance between the win and loss feedbacks was identified between the 190 to 380 ms. Notably, the time window between 330 to 520 ms illustrated a highly significant discrimination effect between IGD and HC groups by using the neural response to loss feedback. In addition, a short-lived group difference between the IGD and HC groups was revealed after perceiving win feedbacks in the time window 330-380 ms.

### 3.7 fNIRS results

To explore where the abnormal corresponding neural response to feedbacks occurred in the IGD group as compared to HCs, the brain activation regions in PFC were identified by using fNIRS neuroimaging.

Pairwise t-tests demonstrated significantly lower activation after win feedbacks in the IGD group was detected as compared to that of the HC group in channel 4 (*t* = −3.003, *p* = 0.0051, covering 36.09% frontopolar area, 42.692% orbitofrontal area, 23.087% inferior prefrontal gyrus), 11 (*t* = −2.94, *p* = 0.0063, covering 29.605% frontopolar area, 70.395% orbitofrontal area), and 13 (*t* = −2.89, *p* = 0.0072, covering 21.429% frontopolar area, 78.571% orbitofrontal area). Meanwhile, significantly lower activation after loss feedbacks was identified in the IGD group as compared to that of the HC group in channel 7 (*t* = - 3.22, *p* = 0.0031, covering 100% frontopolar area) and 14 (*t* = −3.19, *p* = 0.0033, covering 100% frontopolar area).

### 3.8 Correlation results

Our correlation results demonstrated that the risk propensity () was positively correlated with BIS-11 scores for both the HC (*r* =0.47, *p* = 0.013) and IGD groups (*r* =0.61, *p* = 0.016), whereas BIS-11 wa also positively correlated with the number of boxes opened in the IGD group (*r* =0.54, *p* = 0.036) (Fig. 8). No other correlational relations were detected.

**Fig. 6.**
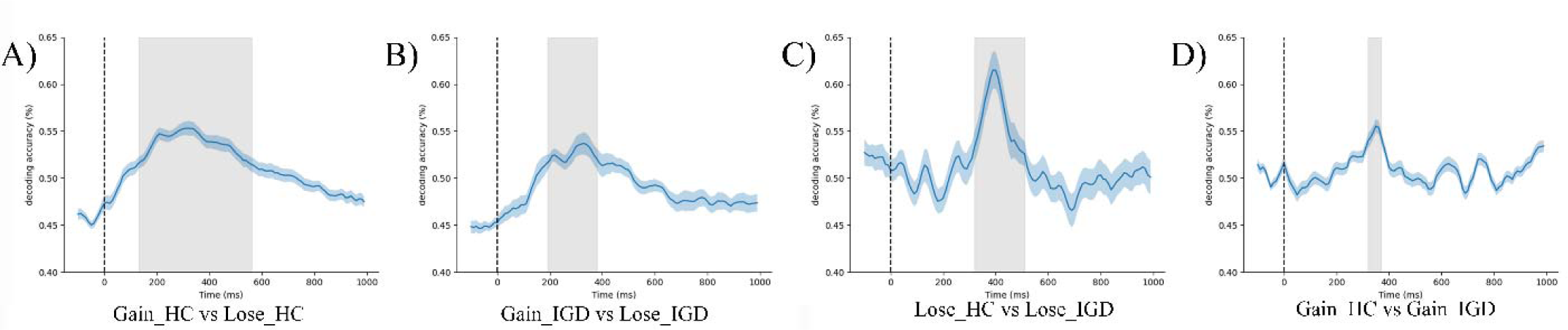
One-vs-one decoding results and neural template analysis (testing for the generalization of decoding across time). A**-D).** Paired wise decoding between the four conditions. Colored curves denoted the decoding accuracy averaged across participants [colored shaded areas: bootstrapped 95% confidence interval; light gray shaded areas: the period of significant decoding at the group level (p < 0.05 lasting for > 20 ms)].

**Fig. 7.**
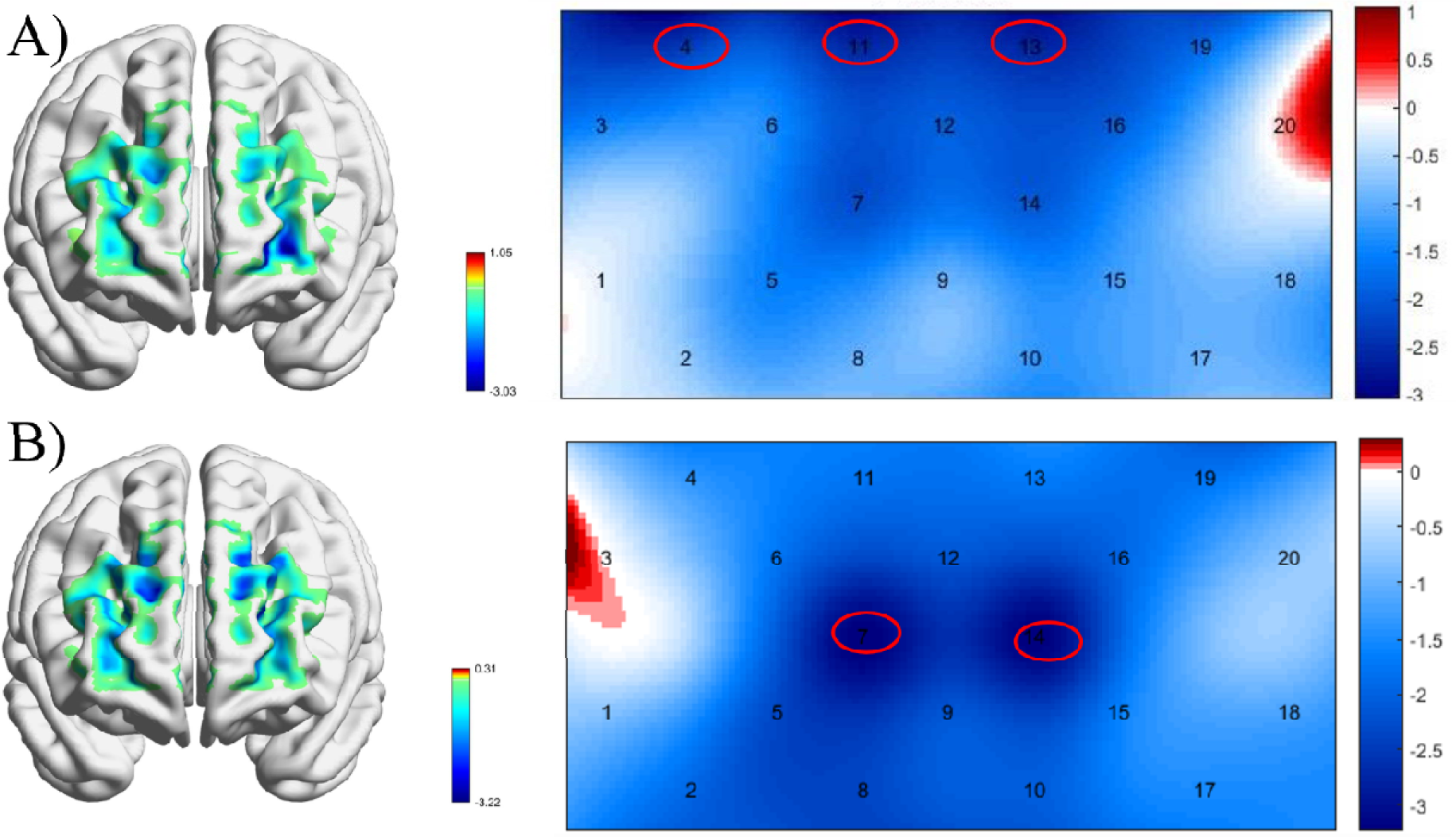
Brain activation difference in PFC between the IGD and HC groups. A). The activation difference after win feedbacks in the IGD group as compared to that of HCs; B) The activation difference after loss feedbacks in the IGD group compared to that of HCs; The color bar denoted the *t* value of contrast.

**Fig. 8.**
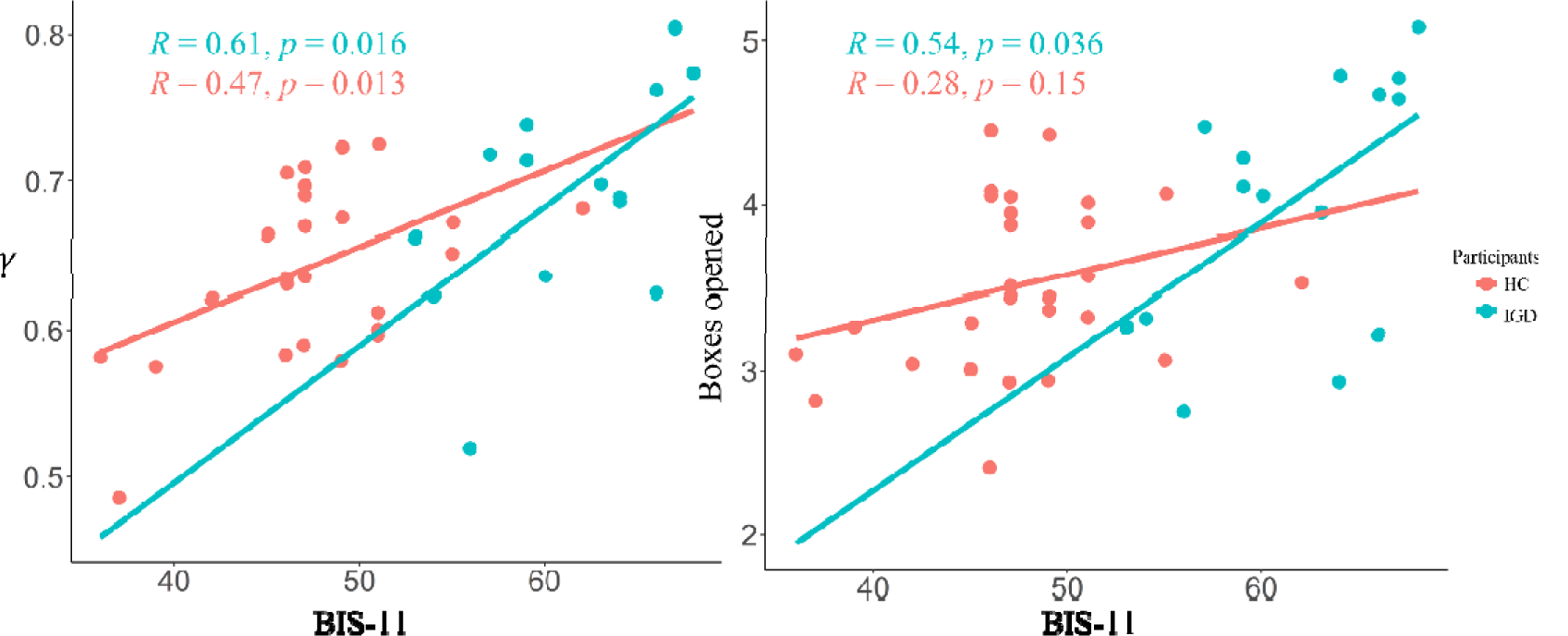
The correlations between demographic data and behavioral data (modelling data for both the IGD and HC groups).

## Discussion

Through the implementation of a sequential risk-taking task and computational modeling, our study revealed that individuals with IGD exhibited a higher propensity for engaging in risky decision-making compared to HCs. Neuroimaging data, including neurophysiological responses P300 and fNIRS assessment of PFC activation, demonstrated distinct neural responses of perceiving outcomes among individuals with IGD compared to HCs. Additionally, our findings have highlighted the impact of outcome valence on subsequent decision-making performance via behavioral results and neuroimaging findings Interestingly, loss outcomes tend to exert a stronger influence on risk-taking propensity compared to win outcomes. Leveraging the power of neuroimaging techniques, we have identified noteworthy variations in neural responses during the P100 and P300 time windows following feedback processing. Furthermore, alterations of theta power brain oscillations in both IGDs and HCs have been observed, particularly in relation to loss outcomes, compared to win outcomes.

First, our behavioral results and modeling parameter findings indicate that individuals with IGD exhibit a heightened inclination towards risk-taking behaviors compared to HCs, which is consistent with previous research on risky propensity of IGD in both ambiguity decision making (Jiang, Li, Zhou, & Zhou, 2020; J. Kim & Kang, 2018) and risky decision making (Ko et al., 2017; Q. Li et al., 2020; L. Liu et al., 2017).

The increased risk propensity observed in individuals with IGD can be attributed to dysregulated reward processing, imbalanced cost-benefit weighting, and poor impulse control. Reward processing refers to the way the brain evaluates and responds to rewarding stimuli, such as pleasurable experiences or outcomes (Meyer, Marco-Pallarés, Boulinguez, & Sescousse, 2021). Previous studies have characterized IGD as an impairment in the reward circuit, with individuals excessively pursuing rewards (L. Wang et al., 2021).

This heightened response to rewards can lead individuals with IGD to be more willing to take risks in pursuit of immediate rewards, often disregarding potential negative consequences or losses (Weinstein & Lejoyeux, 2020). Delay discounting tasks have shown that individuals with IGD are more likely to prioritize immediate rewards (Weinsztok, Brassard, Balodis, Martin, & Amlung, 2021). Additionally, imbalanced cost-benefit weighting, where the expected effect of risky behaviors outweighs the potential loss consequences, contributes to a higher propensity for harmful risk-taking behaviors in individuals with IGD (Hong et al., 2023). While individuals with IGD and HCs may demonstrate similar risk propensities when risky behaviors are favorable, individuals with IGD exhibit significantly higher risk propensities when faced with unfavorable outcomes (Y. W. Yao, Chen, et al., 2015). Lastly, in addition to dysregulated reward processing and imbalanced cost-benefit weighting, poor impulse control might be associated with risky decision making among IGDs. inhibition control plays a crucial role in decision-making to maximize advantages (Sakagami, Pan, & Uttl, 2006). Consequently, previous studies have found that impaired inhibitory control predicted higher addiction degrees and more impulsive in decision making among IGDs (Kräplin et al., 2020; Y. W. Yao, Wang, et al., 2015). The impaired response inhibition and salience attribution (iRISA) model suggests that deficits in inhibition control are core impairments in addiction (Zilverstand, Huang, Alia-Klein, & Goldstein, 2018). Our findings are consistent to this model, as our correlational findings revealed a significantly positive correlation between impulsivity (BIS-11 scores) and risk-taking propensity. As such, impaired inhibition control may be associated with riskier behaviors in sequential risk-taking tasks.

In addition, our study demonstrated significantly different neural responses to feedback among individuals with IGD compared to HCs, as measured by the combined EEG-fNIRS fusion technique. We consistently observed a significant neural response in the P300 time window (300-500 ms) towards loss outcomes in both IGD and HCs using GMD, ERP, and MVPA. The P300 component is a positive-going waveform recorded at centro-parietal sites occurring approximately 300-600 ms after stimulus onset (Yu, Liu, & Shi, 2020), which has been found been associated with processing capacity, mental workload, and also thought to be related to cost-benefit evaluation and decision-making computations (Gui, Li, Li, & Luo, 2016; Kok, 2001). The lower GMD and shorter P300 latency suggested that individuals with IGD made more intuitive decisions in response to feedback. Previous ERP studies have shown that individuals with IGD exhibited reduced P300 amplitudes during auditory information processing, compared to HCs (M. Park, Kim, & Choi, 2016). Also, Y. Zhou, Yao, Fang, and Gao (2022) investigated P300 effects in decision-making processes among individuals with IGD and recreational gaming users but did not find differences between the two groups. There are two possible explanations for this discrepancy. Firstly, our study compared neural responses between individuals with IGD and non-gaming users. Recreational gaming users share more similarity with IGDs in behaviors and brain compared to non-gaming users (Infanti, Valls-Serrano, Perales, Vögele, & Billieux, 2023). Secondly, our study specifically examined neural responses following loss feedback, not the neural response during decision making.

Consistent with previous research, we found reduced activation in the PFC among individuals with IGD after feedback perception, a region known to play important roles in decision-making (St. Onge & Floresco, 2009). Adopting risky decision-making paradigms and neuroimaging tools, Dong and Potenza (2016) observed lower activations in the inferior frontal gyrus while L. Liu et al. (2017) found increased responses in the ventral striatum among individuals with IGD when presented with rewarding outcomes. Higher activation in the reward system may impair the executive control system (Goldstein & Volkow, 2011). Thus, the heightened neural response to feedback among individuals with IGD suggested an enhanced sensitivity to rewards and punishments, indicating dysregulation in the neural circuits involved in reinforcement learning and decision-making processes (Zhang, Hu, Wang, Wang, & Dong, 2020).

Our study found that loss outcomes in decision-making have a significant impact on subsequent decision-making behaviors and elicit distinct neural responses, compared to win outcomes. The influence of feedback on decision-making patterns can be categorized into two distinct patterns: “cold” cognitive patterns and “hot” emotional responses (Y. Zhou et al., 2022). The “cold” cognitive patterns are associated with the central executive network, which evaluates and optimizes decision-making strategies to maximize profit (Salehinejad, Ghanavati, Rashid, & Nitsche, 2021). On the other hand, the feedbacks also trigger emotional and involuntary responses in the body, such as increased heart rate, visceral reactions, perspiration, relief, and regret emotions, which are mediated by the affective network (Weinstein & Lejoyeux, 2020). Our modeling data showed that considering the emotional rating from the previous trial improved the performance of the models, demonstrating that behaviors can be influenced by emotional responses. Loss outcomes tend to elicit stronger emotional responses, which may restrict the involvement of the central executive network (Hsu, Bhatt, Adolphs, Tranel, & Camerer, 2005; Tom, Fox, Trepel, & Poldrack, 2007). Additionally, we observed higher theta oscillations between win and loss feedbacks. This finding is consistent with previous studies indicating that theta oscillations are feedback-related signals and are commonly elicited by incorrect outcomes in these tasks (Y. Wang, Cheung, Yee, & Tse, 2020).

While our study sheds light on the relationship between risky decision-making, neural responses to feedback, and IGD, several limitations should be considered. Firstly, the sample size in our study was relatively small, which may limit the generalizability of the findings. Future research with larger and more diverse samples is necessary to validate and further explore these findings. Additionally, although we excluded patients diagnosed with psychiatric disorders such as anxiety and depression, we did not specifically measure the degree of anxiety and depression among the included participants. This could be taken into consideration in future investigations.

### Conclusions

In summary, our study uncovered that individuals with IGDs exhibit a significant propensity for risk-taking behaviors compared to HCs. This inclination is linked to dysregulated reward processing, poor impulse control, and inhibited inhibitory mechanisms. Neuroimaging results support these findings, showing a heightened sensitivity to rewards and a dearth of cognitive control in IGD. Our insights provide a foundation for future research and potential interventions in the domain of risky decision-making among individuals with IGD.

